# Neural stem cells protect blood-brain barrier integrity via the p38, JNK, and ERK1/2 pathways following intracerebral hemorrhage in rats

**DOI:** 10.1101/2023.09.22.558889

**Authors:** Jing Sun, Xiuli Yang, Austin Carmichael, Tae Jung Kim, Young-Ju Kim, Zhiliang Wei, Ling Han, Nicolas Stanciu, Sang-Bae Ko, Guangxian Nan, Byung-Woo Yoon

## Abstract

Neural stem cells (NSCs) have displayed great potential in ameliorating brain damage following intracerebral hemorrhage (ICH) via proliferation, differentiation, and immunomodulation. However, it remains unclear whether NSCs can improve microvascular function, e.g., blood-brain barrier (BBB) integrity, after ICH. In this study, we investigate the potential therapeutic benefit of NSCs on BBB integrity as well as the underlying mechanism. Adult male Sprague-Dawley rats were randomly divided into sham, ICH+PBS, and ICH+NSCs groups for comparisons. ICH was induced by intrastriatal injection of bacterial collagenase. An aliquot of NSCs or PBS was injected via the tail vein 2 h after ICH induction. The following multiparametric measurements were compared: brain edema, hematoma volume, behavior, BBB permeability, and mitogen-activated protein kinase (MAPK) signaling pathway activity. We found that NSCs treatment attenuates BBB permeability, reduces brain edema, and promotes brain function recovery after ICH by inhibiting ERK1/2, p38, and JNK signaling pathway activation. These findings provide novel insight for future therapies aiming to prevent BBB dysfunction and improve functional recovery in ICH patients.

## 1. Introduction

Despite accounting for only 10-15% of stroke, intracerebral hemorrhage (ICH) deserves attention due to its high rates of morbidity and mortality ^1–5^. ICH is characterized by hematoma formation and expansion in the brain parenchyma following cerebral artery rupture ^1–3^. While primary ICH injury occurs in the first several hours post-ICH as blood components disrupt the brain’s cellular architecture, secondary injury results from the pathologic response to the hematoma over the course of days to weeks ^1–3,5^. Specifically, the activation of pro-inflammatory pathways following microglial activation is thought to play a significant role in ICH secondary brain damage ^3,6,7^. There is also surging evidence suggesting that blood-brain barrier (BBB) disruption contributes to secondary brain injury and worsens neurological outcomes after ICH ^8–11^.

Many studies have implicated the mitogen-activated protein kinase (MAPK) signaling pathway in the regulation of pro-inflammatory cytokines and mediators following ICH ^12,13^. There are several known subfamilies of the MAPK signaling pathway, including the extensively studied and well-characterized extracellular signal-regulated kinase 1/2 (ERK1/2), c-Jun amino terminal kinase (JNK), and p38 MAPK pathways. These pathways are thought to be involved in the regulation of pro-inflammatory cytokines IL-1β and TNF-α as well as the inflammatory mediator COX-2 ^14^. In addition, it has been discovered that inflammatory cytokines can themselves be neurotoxic and exacerbate both BBB permeability and brain edema ^15–18^.

Despite the wide acceptance of advanced surgical decompression to treat ICH, therapeutic options targeting the BBB in acute ICH remain limited. The general benefit of utilizing neural stem cells (NSCs) as a therapy for stroke has been explored in animal models ^19–22^, and it has been found that transplanting NSCs to the injury site improves neurological outcomes via trans-differentiation ^23–25^. Previous studies also indicate that NSCs likely ameliorate ICH-induced secondary brain injury by suppressing the inflammatory response and thus promoting neural cell survival ^23,26^. However, to the best of our knowledge, whether intravenous transplantation of NSCs mitigates BBB disruption after ICH induction remains unclear. Here, we aim to fill this knowledge gap.

In this study, the effects of NSCs on post-ICH brain damage were systematically characterized with emphasis on the BBB integrity and MAPK signaling pathway activity. We hypothesize that NSCs transplantation promotes BBB intactness and ameliorates secondary brain damage by inhibiting both the inflammatory re-sponse and MAPK pathway activation following ICH. To test our hypothesis, we assessed brain edema, hematoma volume, neurological outcome, extravasation of Evans blue, tight junction protein (TJP) expression, inflammatory mediator expression, and MAPK protein phosphorylation in a well-established collagenase-induced ICH rat model.

## 2. Materials and methods

### 2.1. Animals and experimental groups

Our animal study and protocol were approved by the Animal Care and Use of Committee of Seoul National University Hospital (Seoul, South Korea). All animal procedures were performed to minimize pain or discomfort in accordance with current protocols. A total of 78 adult male Sprague-Dawley rats (Orient, Seoul, Republic of Korea) aged 8 weeks were used in the study under the experimental paradigm outlined in Figure 1. Animals were housed under a 12 h light/dark cycle with free access to food and water. These rats were randomly assigned to three experimental groups, i.e., one control group (denoted as sham) and two experimental groups receiving phosphate buffer saline (PBS) and NSCs treatments (denoted as ICH+PBS and ICH+NSCs, respectively).

**Figure 1.**
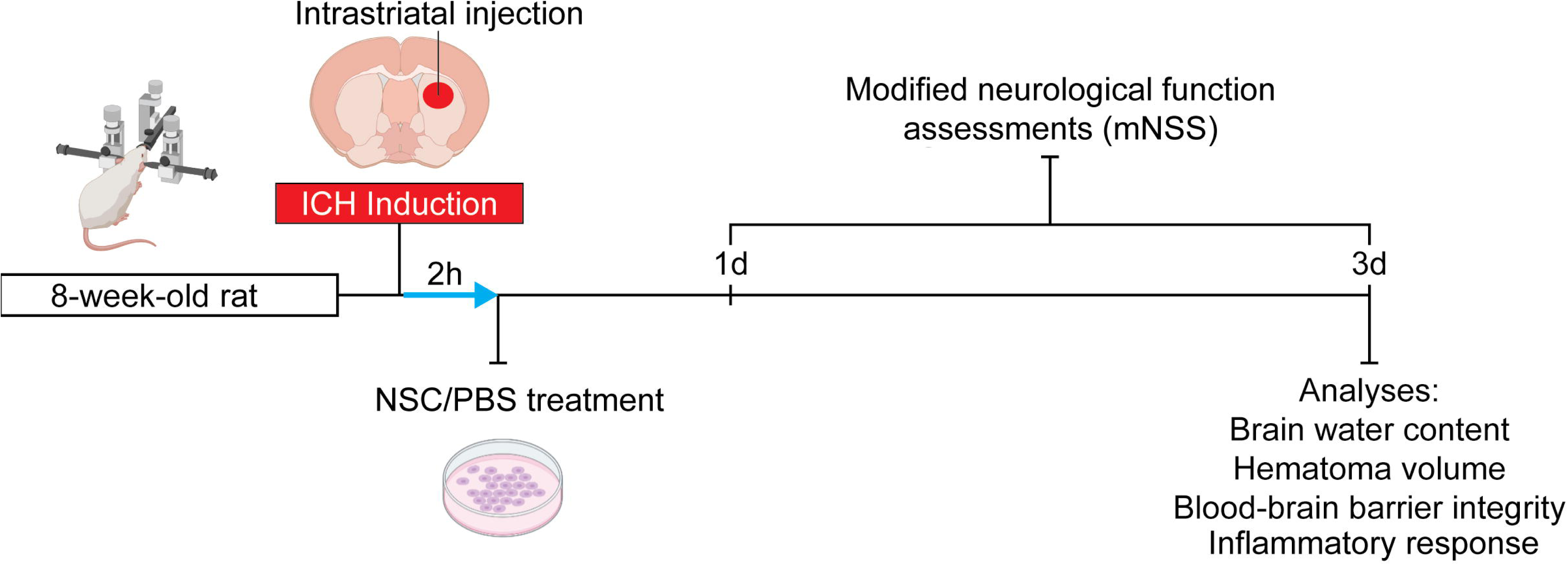
Schematic representation of experiments.

### 2.2. Establishment of the ICH model and cell transplantation

ICH was induced by the stereotaxic injection of collagenase type IV (Sigma-Aldrich, USA) following the reported schemes with some modifications ^27,28^. After anesthetic induction with 2% isoflurane delivered by 70% N_2_/30% O_2_, rats were placed in the stereotaxic frame. Under aseptic conditions, an incision was performed along the sagittal midline of rat head to expose the bregma. A burr hole was made using the coordinates of 2.0 mm anterior, 6.0 mm ventral, and 3.0 mm lateral to the bregma. A Hamilton syringe was inserted stereotactically through the burr hole and into the left striatum. Collagenase IV (0.4 U) in 0.8 μl saline was injected over a period of 5 min. After holding placement for another 2 min, the microsyringe was slowly removed. The burr hole was then sealed with bone wax, and the skin incision was sutured. Rats of the sham group were manipulated identically except that 0.8 μl sterile saline was administered instead of collagenase. Rectal temperature was monitored and maintained at 37.0 ± 0.2°C throughout the surgical procedures using a homeothermic blanket system (Harvard Apparatus, Holliston, USA).

HB1.F3 cells, a stable immortal human neural stem cell line ^25^, were administered via the tail vein 2 h post-ICH (4×10^6^ cells in 0.5 ml PBS). As a comparison, an equal volume of PBS without cells was administered in ICH+PBS group.

### 2.3. Brain water content measurement

Brain water content was assessed to investigate post-ICH edema. At 72 h post-ICH, brains were harvested and divided into two hemispheres along the midline after the cerebellum and brain stem were removed. Brain tissues were weighed on an electronic analytical balance to obtain the wet mass (denoted as *M_wet_*) and then dried in a 100°C oven for 24 h to obtain the dry mass (denoted as *M_dry_*). The brain water content was calculated using the following formula: 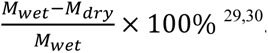 ^29,30^.

### 2.4. Hematoma volume measurement

Rat brains were harvested for morphometric analysis at 72 h post-ICH induction. Rat brains were cut coronally through the needle entry plane and then serially sliced (1-mm thickness) anteriorly and posteriorly by respect to the needle entry site. Digital photographs of the serial slices were taken, and the hematoma volume was calculated using an image analysis program (ImageJ, National Institutes of Health, Baltimore, USA). The total hematoma volume (mm^3^) was calculated by multiplying the hemorrhagic area in each section by the distance between sections ^31^.

### 2.5. Neurological function assessments

Neurological outcomes were rated using the modified Neurological Severity Score test (mNSS) at 24 h and 72 h post-ICH. The mNSS is a composite of motor (muscle status and abnormal movement), sensory (visual, tactile, and proprioceptive), balance, and reflex tests. Neurologic function was graded on a scale of 0-18 (normal score: 0; maximal deficit score: 18) ^32,33^. A lower mNSS score indicates better neurological recovery. Rats were trained and assessed prior to surgery to ensure that basal scores were 0. All behavioral tests were conducted in a quiet and dimly lit room by two experimenters blinded to the group identifications.

### 2.6. Blood-brain barrier integrity by Evans blue extravasation

At 72 h after ICH induction, 4.0% Evans blue (EB, Sigma, USA) was intravenously administered via the tail vein at the injection volume of 2µl/g ^34^. Two hours after the Evans blue injection, rats were anesthetized and underwent transcardial perfusion with cold saline until the fluid from the right atrium appeared colorless. Then, the brain was rapidly harvested and cut into two hemispheres. The brain tissue was weighed and mechanically homogenized in 3 mL formamide (Sigma, USA). The obtained suspension was kept at 60°C in a water bath in the dark for 24 h and then centrifuged at 12,000×g for 30 min. The supernatant was collected and spectrophotometrically analyzed (λ = 620 nm) with a microplate reader (Molecular Devices, VersaMax ELISA microplate reader, USA) to determine the Evans blue content. Evans blue extravasation into the brain parenchyma was assessed with a standard curve and expressed as μg/g of wet brain tissue.

### 2.7. Tissue preparation and immunofluorescence staining

For the immunofluorescence analysis of the perihematomal tissue ^35^, rats were anesthetized and underwent transcardial perfusion with 100 mL of saline followed by 100 mL of 4% paraformaldehyde (Merck, Germany) at 72 h following ICH induction. After the brain samples were fixed in 4% paraformaldehyde overnight, they were dehydrated in 30% sucrose (Daejung, Korea) and embedded in frozen section compound (Leica, Germany). Coronal sections (10 µm) were prepared using a freezing microtome (Leica, German). Before the sections were stained, they were washed with PBS three times and blocked with blocking buffer (0.5% BSA, 3% Triton-100, and 10% normal goat serum) for 1 h at room temperature. Then, the sections were incubated overnight at 4°C with the following primary antibodies: mouse anti-claudin-5 (Invitrogen, USA, 1:100 dilution) and rabbit anti-occludin (Invitrogen, USA, 1:100 dilution). The sections were washed three times in PBS and were incubated in the dark for 1 h at room temperature with goat anti-mouse Alexa Fluor^®^ 488-conjugated antibody (Invitrogen, USA, 1:200 dilution) or goat anti-rabbit Alexa Fluor^®^ 488-conjugated antibody (Invitrogen, USA, 1:200 dilution). The sections were then washed and covered with mounting medium containing 4’6’-diamino-2-phenylindole (DAPI) (Vector, USA). Images of the sections were acquired using a fluorescence microscope (Leica, DM5500B, German).

### 2.8. Western blot assay

The Western blot analysis was performed as described in previous studies ^36–38^. The peri-hemorrhage brain samples were homogenized, and the protein was extracted using radio immunoprecipitation assay lysis buffer. The homogenate was incubated for 30 min on ice and then centrifuged at 14,000×g for 30 min at 4°C. Protein concentrations were determined using a Micro Bradford Protein Assay Kit (Bio-Rad, Rockford, IL, USA). Samples with equal quantities of protein (50 μg) were separated on 10% or 15% sodium dodecyl sulfate polyacrylamide gels, transferred onto polyvinylidene difluoride (PVDF) membranes, and blocked in 5% skim milk buffer at room temperature for 1 h. The membranes were then incubated overnight at 4°C with the following primary antibodies: anti-COX-2 (Abcam, UK, 1:1000 dilution), anti-IL-1β (Abcam, 1:1000 dilution), anti-occludin (Invitrogen, USA, 1:1000 dilution), anti-claudin-5 (Invitrogen, USA, 1:1000 dilution), anti-p-ERK1/2 (Cell Signaling, USA, 1:500 dilution), anti-p-p38 (Cell Signaling, USA, 1:500 dilution), anti-p-JNK (Cell Signaling, USA, 1:500 dilution), anti-ERK1/2 (Cell Signaling, USA, 1:1000 dilution), anti-p38 (Cell Signaling, USA, 1:1000 dilution) and anti-JNK (Cell Signaling, USA, 1:1000 dilution). Anti-β tubulin (Bioss, USA, 1:10,000 dilution) was used as the loading control. Blot bands were quantified using densitometry with ImageJ software (National Institutes of Health, Baltimore, MD, USA). Phosphorylation level of proteins was indicated by the ratio of phosphoprotein to total protein. The value was then normalized to the mean value of the sham group for comparison.

### 2.9. Enzyme-linked immunosorbent assay (ELISA)

The rat brain tissue was harvested at 72 h after ICH induction or sham operation. The levels of TNF-α in tissues were quantified by ELISA ^39^. Photometric measurements were conducted at 450 nm using a microplate reader (Molecular Devices, Versa Max ELISA microplate reader, USA). For the ELISA process, commercial ELISA kits (R&D Systems, USA) were used following the manufacturer’s instructions. The level of TNF-α was expressed as pg/mL.

### 2.10. Data processing and statistical analysis

Quantitative analyses were performed by investigators who were blinded to the experimental protocol and animal identity. All data are shown as the mean ± standard error of the mean (SEM) and were statistically analyzed using one-way analysis of variance (ANOVA) followed by Tukey’s post hoc test with multiple comparisons, except for behavioral tests and hematoma volume, which were analyzed by Mann-Whitney U test. *P* < 0.05 was considered a statistically significant difference. All statistical analyses were performed using GraphPad Prism version 10.0.

## 3. Results

### 3.1. NSCs treatment decreases brain water content and improves functional recovery after ICH

Brain water content was assessed to determine the effect of NSCs transplantation on brain edema post-ICH. At 72 h post-operation, the ICH+NSCs group displayed marginally lower brain water content compared to the ICH+PBS group (Fig. 2a, *P* < 0.05; *F* = 38.40). This suggests that NSCs treatment reduces brain edema. Hematoma volumes were similar between the ICH+PBS and ICH+NSCs groups at 72 h post-operation (Fig. 2b, *P* > 0.05).

**Figure 2.**
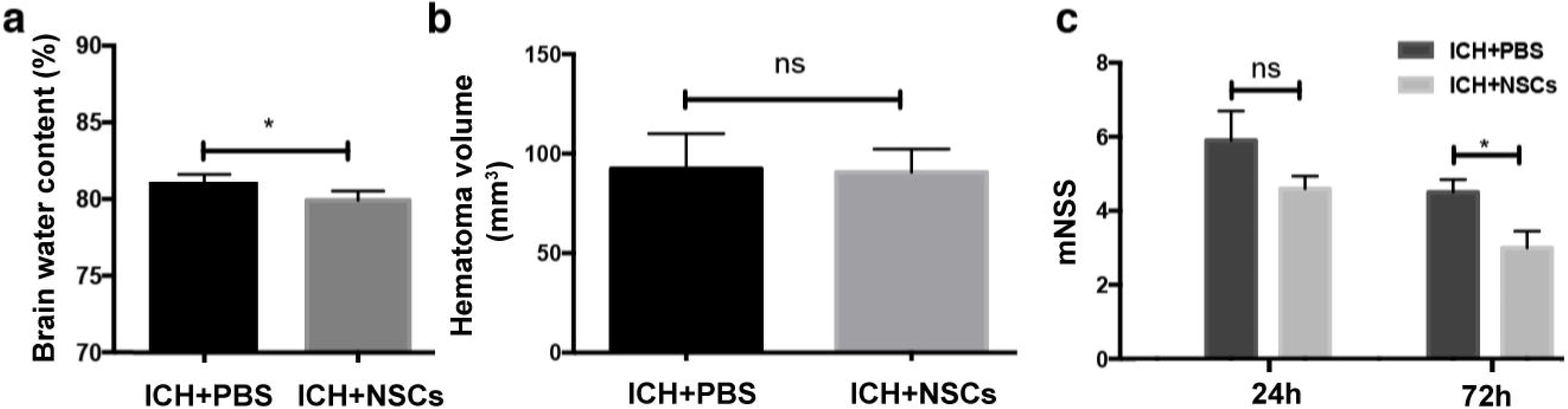
The effect of NSCs treatment on brain water content, hematoma volume, and neurological recovery following ICH. (**a**) At 72 h post-ICH induction, the brain water content of the ICH+NSCs group was marginally lower than the ICH+PBS group. (**b**) Hematoma volume did not differ significantly between the ICH+PBS and ICH+NSCs groups. (**c**) All rats were subjected to modified Neurologic Severity Score (mNSS) assessments. At 24 h post-ICH induction, there was no significant difference in neurological recovery between the ICH+PBS and ICH+NSCs groups at 24 h post-ICH induction. At 72 h post-operation, the ICH+NSCs group displayed lower mNSS scores and thus improved neurological recovery relative to the ICH+PBS group. Data are represented as mean ± SEM. ns = not significant, \**P* < 0.05 by one-way ANOVA with Turkey’s post-hoc test (n=8).

mNSS tests were also performed to assess the effect of NSCs treatment on neurological function at 24 h and 72 h post-ICH inducement. No differences in neurological outcome were observed at 24 h. However, NSCs treatment significantly reduced mNSS scores and thus improved neurological recovery at 72 h post-operation (Fig. 2c, *P* < 0.05).

### 3.2. NSCs treatment decreases BBB permeability and increases TJPs expression following ICH

BBB disruption and edema formation are associated with endothelial dysfunction. We investigated the neurovascular protective action of NSCs in ICH by assessing Evans blue extravasation 72 h post-ICH induction (Fig. 3a). The ICH+PBS group exhibited significantly higher extravasation of Evans blue (Fig. 3b, *P* < 0.05) compared to the sham group, indicating post-ICH BBB damage. In contrast, the ICH+NSCs group was associated with significantly less dye extravasation relative to the ICH+PBS group (Fig. 3b, *P* < 0.05, *F* = 8.046), indicating that NSCs treatment has a BBB-protective effect.

**Figure 3.**
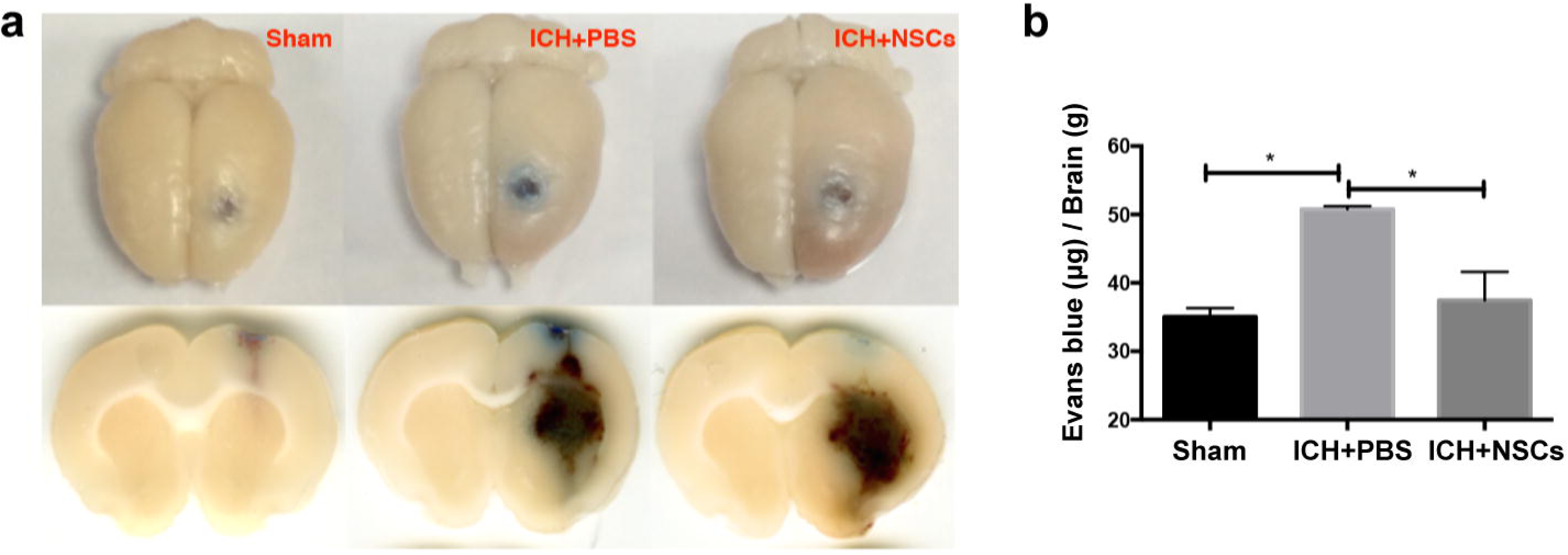
NSCs treatment reduces Evans blue (EB) extravasation and BBB permeability. (**a**) Images depict EB leakage into the brain parenchyma for the sham, ICH+PBS, and ICH+NSCs groups at 72 h post-operation. (**b**) Greater EB extravasation occurred in the ICH group compared to the sham group, while the ICH+NSCs group displayed less EB leakage relative to the ICH+PBS group at 72 h post-operation. Data are represented as mean ± SEM. \**P* < 0.05 by one-way ANOVA with Turkey’s post-hoc test (n=8).

We further investigated the microvascular underpinning of BBB integrity by monitoring the expression of two TJPs: occludin and claudin-5. The ICH+PBS group was associated with reduced occludin expression relative to the sham group. Based on the immunofluorescent staining (Fig. 4a) and western blot (Fig. 4b and c) analyses, NSCs treatment ameliorated occludin losses (Fig. 4, *P* < 0.05, F=30.68) following ICH. The immunofluorescence (Fig. 5a) and western blot analyses of claudin-5 showed similar results as occludin (Fig. 5b and c, *P* < 0.05, *F* = 51.73). Together, these findings reveal that NSCs treatment may bolster BBB stability by salvaging TJP expression and activity following ICH.

**Figure 4.**
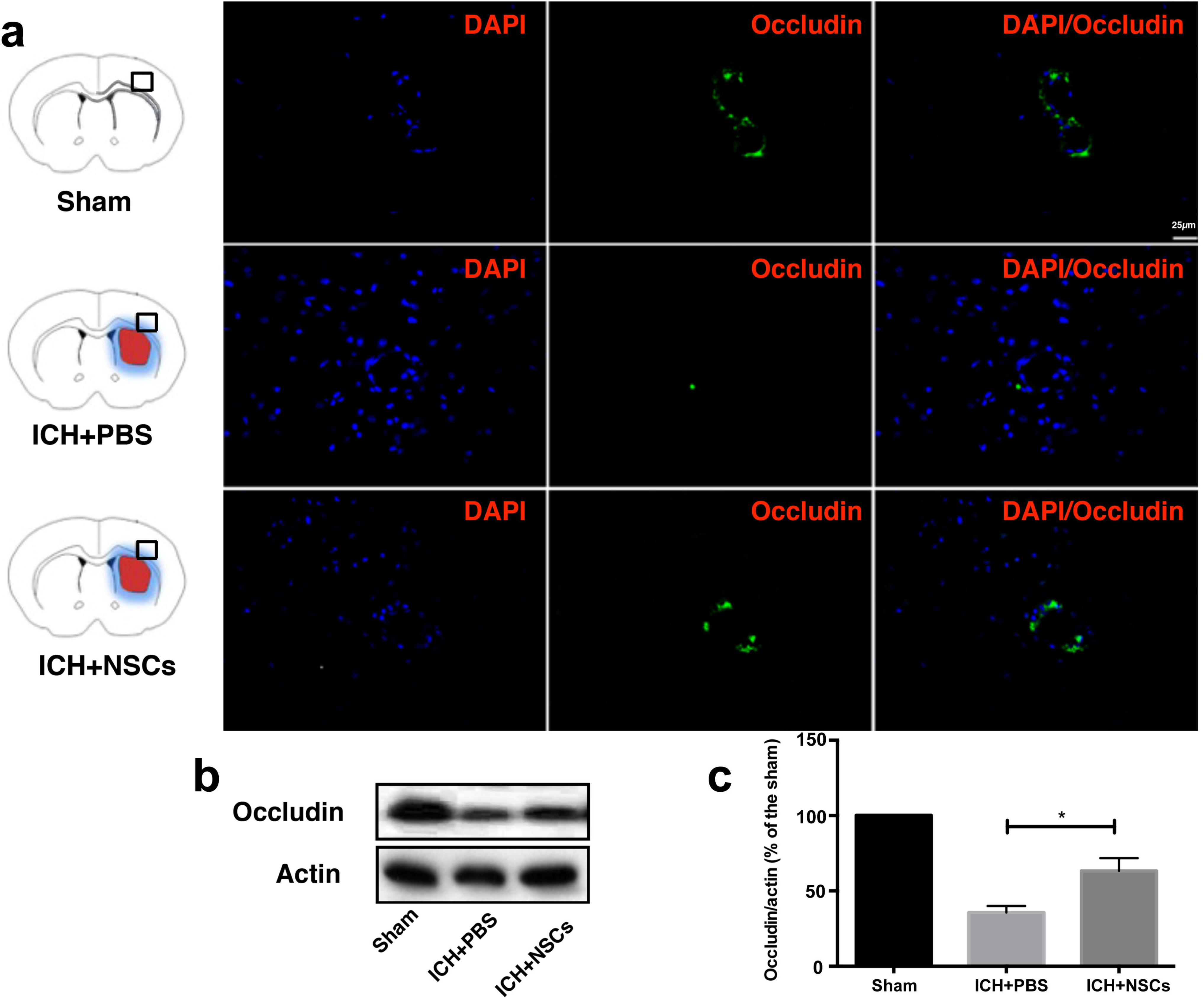
NSCs treatment rescues the expression of tight junction protein occludin. (**a**) Representative microphotographs showed that NSCs treatment partially recovered occludin expression at 72 h post-ICH induction. Scale bar = 25 µm. (**b, c**) At 72 h post-operation, western blot analysis revealed a lower density of occludin in the ICH groups relative to the sham group. Compared to the ICH+PBS group, the ICH+NSCs group displayed greater occludin density, suggesting that NSCs treatment leads to a partial recovery in occludin expression following ICH. Data are represented as mean ± SEM. \**P* < 0.05 by one-way ANOVA with Turkey’s post-hoc test (n=5).

**Figure 5.**
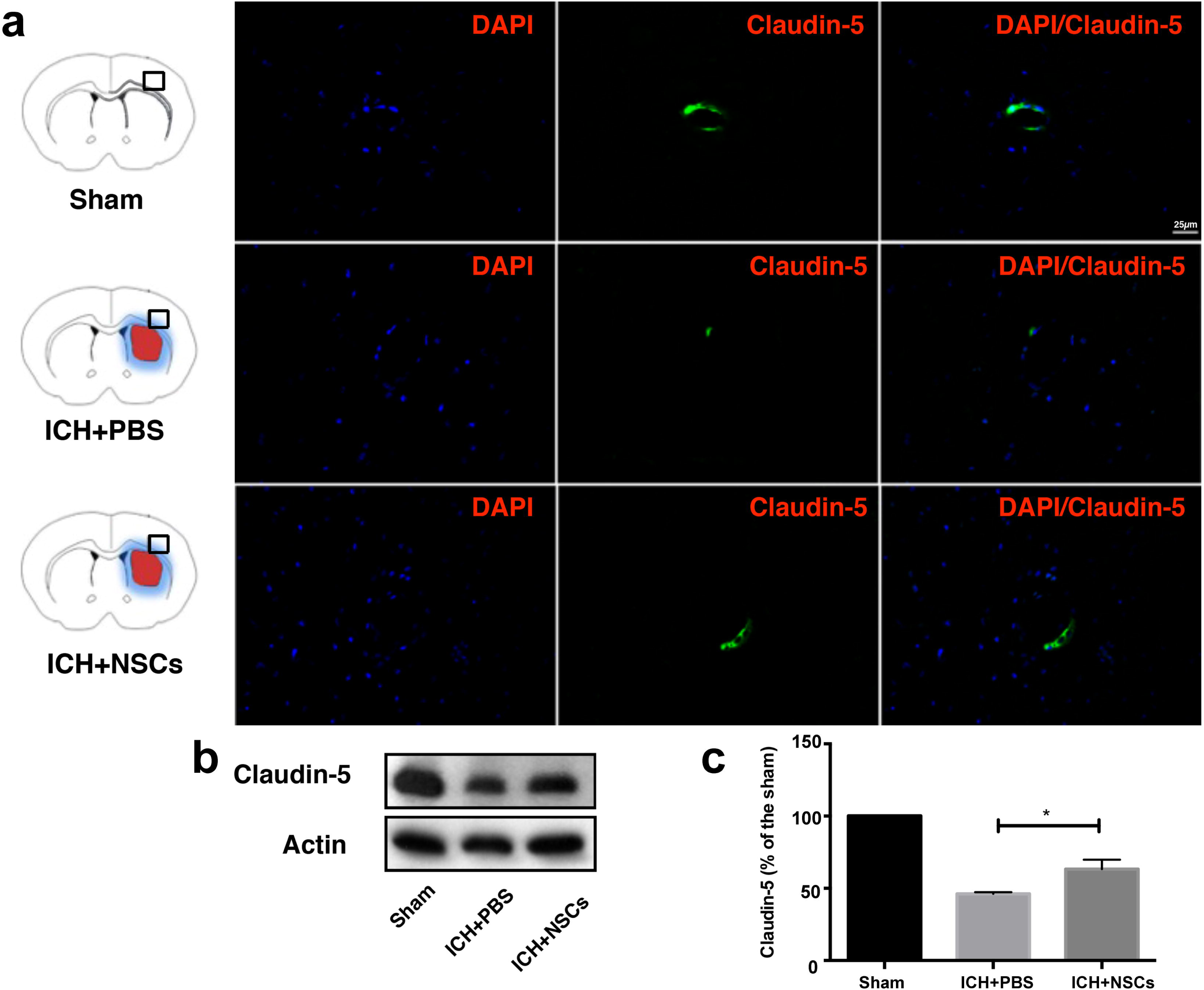
NSCs treatment rescues the expression of tight junction protein claudin-5. (**a**) Representative microphotographs showed that NSCs treatment partially recovered claudin-5 expression at 72 h post-ICH induction. Scale bar = 25 µm. (**b, c**) At 72 h post-operation, western blot analysis revealed a lower density of claudin-5 in the ICH groups relative to the sham group. Compared to the ICH+PBS group, the ICH+NSCs group displayed greater claudin-5 density, suggesting that NSCs treatment leads to a partial recovery in claudin-5 expression following ICH. Data are represented as mean ± SEM. \**P* < 0.05 by one-way ANOVA with Turkey’s post-hoc test (n=5).

### 3.3. NSCs treatment reduces inflammatory mediator expression and the phosphorylation of MAPK signaling pathway-related proteins

To further assess brain microenvironments, which may be closely related to BBB breakdown, we examined the expression of inflammatory mediators and MAPK signaling pathway-related proteins in the injury lesion at 72 h post-ICH induction (Fig. 6a-h). The expression levels of inflammatory mediators COX-2, IL-1β, and TNF-α were increased in the ICH+PBS and ICH+NSCs groups compared to the sham group. Relative to PBS treatment, NSCs treatment significantly reduced the expression of inflammatory mediators COX-2 (Fig. 6b, *P* < 0.01; *F* = 48.78), IL-1β (Fig. 6c, *P* < 0.05; *F* = 19.93), and TNF-α (Fig. 6d, *P* < 0.05; *F* = 9.030).

**Figure 6.**
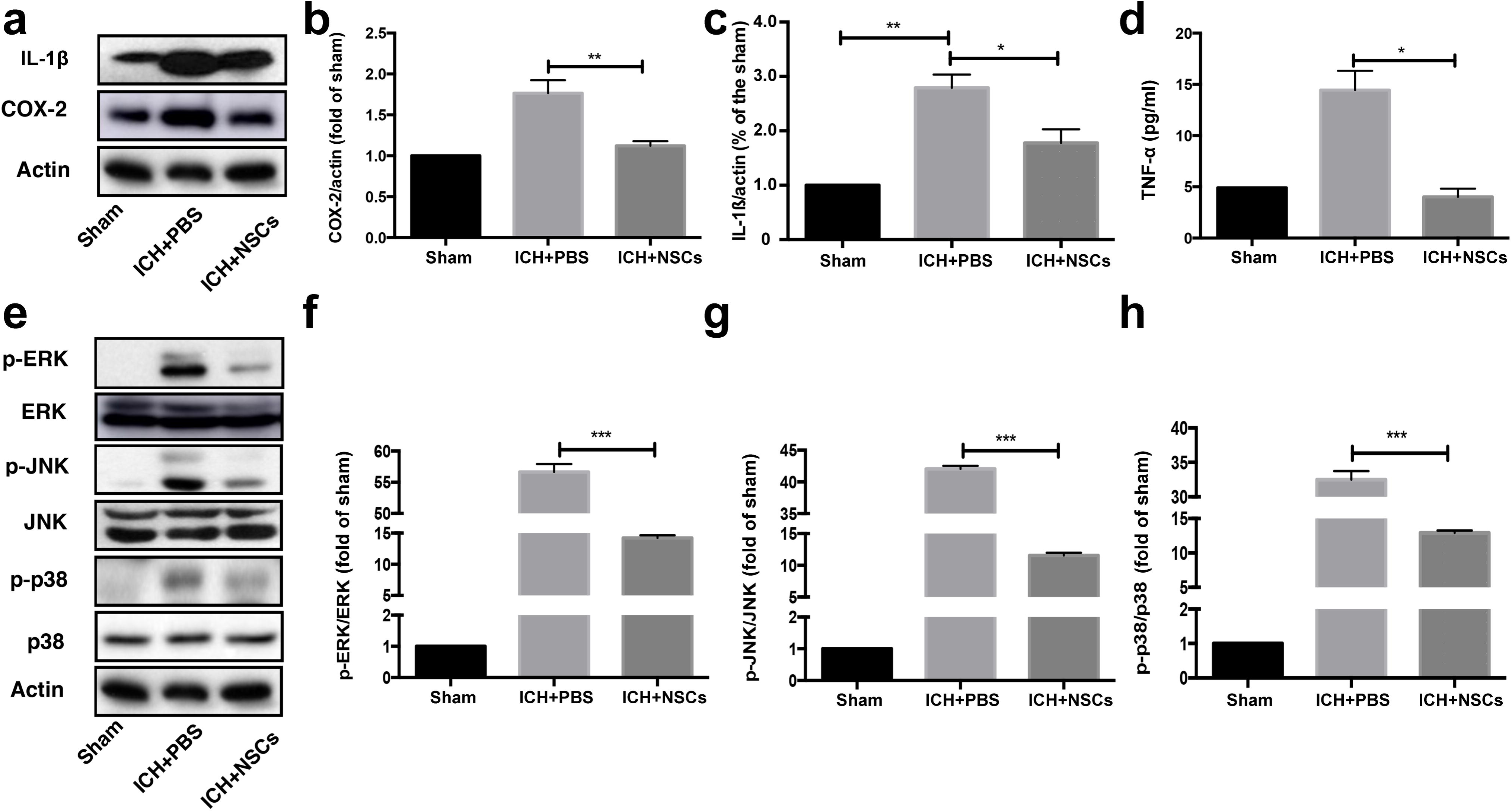
NSCs treatment attenuates inflammatory mediator expression and MAPK signaling pathway activation. (**a-d**) At 72 h post-operation, the ICH+PBS group displayed a greater density of COX-2 (72 kDa) and IL-1β (31 kDa) as well as an increased level of TNF-α in the brain relative to the sham group. (**e-h**) The ICH+PBS group had a greater ratio of phosphorylated for ERK (42 and 44 kDa), JNK (46 and 54 kDa), and p38 (43 kDa) MAPK proteins relative to the sham group at 72 h post-ICH induction. NSCs treatment attenuated the ICH-induced phosphorylation of MAPK signaling pathways. Data are represented as mean ± SEM. \**P* < 0.05, \**P* < 0.01, and \*\*\**P* < 0.001 by one-way ANOVA with Turkey’s post-hoc test (n=5).

Compared with basal levels in the sham group, the phosphorylation levels of ERK1/2, JNK, and p38 MAPK proteins robustly increased after ICH inducement. Relative to PBS treatment, NSCs treatment inhibited the phosphorylation of ERK1/2, JNK, and p38 (Fig. 6e-h, *P* < 0.001). These findings suggest that NSCs reduce MAPK signaling pathway activity following ICH injury.

## 4. Discussion

In this study, we present the potential therapeutic benefit of NSCs in attenuating BBB disruption during the early stage of ICH recovery. We found that NSCs transplantation mitigates BBB permeability and promotes functional recovery in rats after ICH induction. Regarding underlying molecular mechanisms, we found that NSCs treatment reduces the inflammatory response and suppresses p38, ERK1/2, and JNK signaling pathway activation.

Complementary to existing studies on the therapeutic benefit of NSCs in animal stroke models, this study associates functional recovery with BBB intactness ^23–25^. Numerous studies have demonstrated that BBB disruption plays a critical role in post-stroke brain injury and correlates with poor outcomes ^37,40,41^. It has also been established that disrupting the components of the BBB prompts brain edema formation and disease deterioration, which is consistent with our finding of lower TJP expression and greater Evans blue extravasation in the ICH+PBS group. Building upon these principles, our study reveals that early intravenous delivery of NSCs restores the expression of TJPs, reduces brain edema, promotes BBB integrity, and enhances neurological recovery after ICH. Although further explorations are required to establish a concrete mechanism of action, NSCs may salvage BBB impermeability through antioxidative effects, by enhancing the survival of neural cells, and reducing inflammatory cell infiltration. These findings demonstrate the therapeutic promise of NSCs in maintaining the structural composition of the BBB following ICH.

To better understand the mechanism by which NSCs treatment reduces BBB permeability and promotes functional recovery following ICH, we explored the effect of NSCs on inflammatory mediator expression and MAPK signaling pathway activity. Inflammatory responses, including the infiltration of neutrophils and mononuclear macrophages as well as the activation of microglia and astrocytes, are known to play a critical role in ICH-induced secondary brain injury ^42–44^. It has been demonstrated that the excessive upregulation of inflammatory mediators, including IL-1β, TNF-α, and COX-2, in the perihematomal region after ICH is strongly correlated with brain edema, BBB dysfunction, and neuronal ferroptosis ^36^. Additionally, MAPK signaling pathway activation has been implicated in both the regulation of pro-inflammatory cytokines as well as BBB destabilization via TJP downregulation. In our study, the expression of these cytokines significantly increased at 72 h post-ICH, which is consistent with the timing of increased brain edema and BBB disruption following ICH ^45^. Interestingly, NSCs treatment lowered inflammatory mediator expression, suggesting that NSCs may promote neurological recovery after ICH by hindering the inflammatory response. Likewise, NSCs treatment reduces the phosphorylation of MAPK signaling pathway-related proteins. Together, these findings suggest that the mechanism by which NSCs promote BBB stability and neurological recovery after ICH likely entails reducing the inflammatory response by inhibiting MAPK signaling pathway activity.

While we are optimistic that our findings may provide valuable insight for future therapies, our project is not without limitations. First, it has been reported that BBB disruption occurs as early as 0.5 h following ICH induction, reaches a peak at 72 h, and declines to a normal state by 2 weeks post-ICH. In this study, we investigated the effect of NSCs transplantation on BBB integrity at 24 h and 72 h post-ICH. Thus, we did not observe the dynamic effect of NSCs on the BBB integrity at the most-acute stage post-ICH. It is our plan to utilize advanced magnetic resonance imaging methods to monitor the dynamic events associated with microvascular function in the acute stage of ICH ^46–49^. Second, our study focused on the effect of NSCs on TJPs in the BBB. Other microvascular structural components, such as astrocytic end-feet and pericytes, are also pertinent to BBB integrity and may be impacted by NSCs treatment ^50^. Third, there are several alternative mechanisms by which NSCs could impact BBB permeability that were not explored, including via antioxidative effects. Further studies will be required to verify other potential mechanisms.

## 5. Conclusion

Through a well-established rat model, we demonstrated that NSCs treatment rescues BBB integrity and improves functional outcomes in the early stage of ICH recovery. We found that NSCs decrease the inflammatory response via modulating the MAPK pathway activity following ICH. Therefore, we propose that the ERK1/2, JNK, and p38 signaling pathways may play key roles in the mechanism by which ICH induces secondary brain injury. We are optimistic that future studies may reveal NSCs transplantation as a viable ICH treatment strategy.

## Acknowledgments

This research was supported by grants from the Ministry of Health and Welfare, Republic of Korea (Grant No. HI15C2311).

## Competing Interests

The authors declare no competing interests.

## Compliance with ethical standards

All animal experiments and procedures in this study were performed in accordance with the ethical standards described in the methods section.

## Notes

### Competing Interest Statement

The authors have declared no competing interest.

### Summary of Updates

Minor revisions to improve clarity. Authorship updated.

